# RNA-dependent aggregation of a common TEV protease variant alters *in vitro* biomolecular condensate reconstitution

**DOI:** 10.64898/2026.06.16.732646

**Authors:** Joseph A Larson, Devyn Iglesias-Fuller, Andrea A Putnam

## Abstract

Biomolecular condensates formed by liquid-liquid phase separation (LLPS) are commonly studied *in vitro* using protease-mediated removal of solubilizing tags to induce condensation under controlled conditions. Tobacco Etch Virus (TEV) protease is widely used for this purpose and is generally assumed to remain soluble and inert during condensate reconstitution. Here, we show that in RNA-containing systems, TEV protease variants can interact with RNA, leading to aggregation and changes in the phase behavior of the target protein. Using confocal microscopy, turbidity measurements, and mass photometry, we demonstrate that commonly used TEV protease variants differ in their propensity to undergo RNA-dependent aggregation. The widely used pRK793 TEV protease forms large RNA-associated aggregates. We further show that RNA-TEV aggregation alters the morphology and organization of protein-RNA condensates formed by well-characterized phase-separating proteins, including PGL-3 and FUS. Together, our findings show that TEV protease can directly impact *in vitro* LLPS assays through RNA binding and aggregation. These results underscore the importance of validating protease-based induction strategies and incorporating appropriate controls when reconstituting biomolecular condensates, particularly in RNA-rich systems.

## Introduction

Liquid-liquid phase separation (LLPS) occurs when macromolecules demix from a homogeneous solution to form dense and dilute liquid phases. In cells, LLPS has been proposed to underlie the formation of membraneless compartments known as biomolecular condensates; assemblies enriched in proteins and nucleic acids. A widely used strategy to assess whether a protein undergoes LLPS is purification followed by *in vitro* condensate reconstitution, often in the presence of RNA (Lin et al. 2015; Wadsworth et al. 2024).

A recurring challenge in the study of condensates is that proteins prone to phase separation frequently exhibit limited solubility when expressed recombinantly, complicating purification and biochemical characterization (Alberti et al. 2019; Putnam and Seydoux 2021). To overcome this limitation, target proteins are commonly fused to solubilizing tags that suppress premature condensation during expression and purification. These tags are typically removed proteolytically immediately prior to or during *in vitro* assays, enabling temporally controlled induction of phase separation without altering buffer composition, ionic strength, or denaturant concentration (Burke et al. 2015; Rhine et al. 2020; Currie et al. 2023). Tobacco Etch Virus (TEV) protease is one of the most commonly used enzymes for this purpose.

TEV protease is a cysteine protease that cleaves a seven-amino acid recognition sequence (Parks et al. 1994). Its exceptional sequence specificity and minimal off-target cleavage have made TEV protease a standard tool for removing solubility or affinity tags from recombinant proteins for decades (Terpe 2003). However, native TEV protease exhibits limited solubility and undergoes auto-inactivation through self-cleavage (Kapust et al. 2001). To address these limitations, multiple engineered TEV variants have been developed that improve solubility and stability, including the incorporation of affinity or epitope tags, point mutations, and elimination of the self-cleavage site (Kapust et al. 2001; Nautiyal and Kuroda 2018). A widely used variant includes an N-terminal polyhistidine tag for immobilized metal affinity chromatography, a C-terminal arginine-rich tag, and the S219V mutation to prevent auto-cleavage and improve catalytic efficiency, yielding a ∼28 kDa protease commonly used in biochemical and biophysical assays (Tropea et al. 2009).

Because TEV protease is routinely assumed to remain soluble and inert under cleavage conditions, its potential contribution to phase separation has received limited attention. However, early biochemical studies reported that TEV protease can interact with RNA, but these observations have not been widely cited or incorporated into current experimental practice (Daròs and Carrington 1997). Moreover, the engineered features of TEV protease, including arginine-rich regions, raise the possibility that TEV variants may engage in electrostatic interactions with RNA, particularly under conditions commonly used for condensate reconstitution.

Here, we show that several commonly used TEV protease variants interact with RNA and undergo RNA-dependent aggregation *in vitro*. We find that the widely used pRK793 TEV protease exhibits aggregation behavior that depends on RNA identity and concentration, and that this behavior can alter the morphology and organization of protein-RNA condensates. These findings demonstrate that TEV protease is not universally inert in LLPS assays and can influence condensate properties through RNA-mediated interactions. We conclude by proposing best practices for the use of TEV protease in *in vitro* condensate experiments and by outlining experimental controls that can mitigate unintended effects.

## Results

### RNA aggregation by TEV protease variants

To test whether commonly used TEV protease variants aggregate with RNA, we examined aggregation in the presence of the long single-stranded RNA *nos-2* (1248 nt), which has been previously used to induce phase separation of the protein PGL-3 (Putnam et al. 2019; Folkmann et al. 2021). We analyzed four widely used TEV protease variants: **pRK793**, which contains the S219V mutation to reduce autocleavage, an N-terminal 7×His tag for purification, and a C-terminal 5×Arg tag to enhance solubility (Kapust et al. 2001); **His::TEV**, a pRK793-derived variant lacking the C-terminal 5×Arg tag; **SuperTEV**, a pRK793-derived construct lacking the 5×Arg tag and containing nine additional mutations designed to improve solubility (Lau et al. 2023); and **AcTEV**, a commercially available TEV protease preparation (Supplemental Table 1). To control for preparation-specific effects, we also analyzed an independently prepared pRK793 TEV protease sample (**pRK793 TEV-2**).

We first assessed RNA aggregation by confocal imaging of fluorescently labeled *nos-2* RNA. In the absence of protease, RNA was uniformly distributed (Fig. 1A). Addition of either pRK793 TEV or pRK793 TEV-2 resulted in the formation of prominent RNA aggregates (Fig. 1A), indicating that aggregation was not specific to a single protein preparation. SuperTEV protease also induced RNA aggregation, although to a lesser extent than pRK793 TEV. In contrast, no observable RNA aggregates were detected following addition of AcTEV or His::TEV protease (Fig. 1A), demonstrating that RNA aggregation is strongly dependent on the TEV variant used.

**Figure 1:**
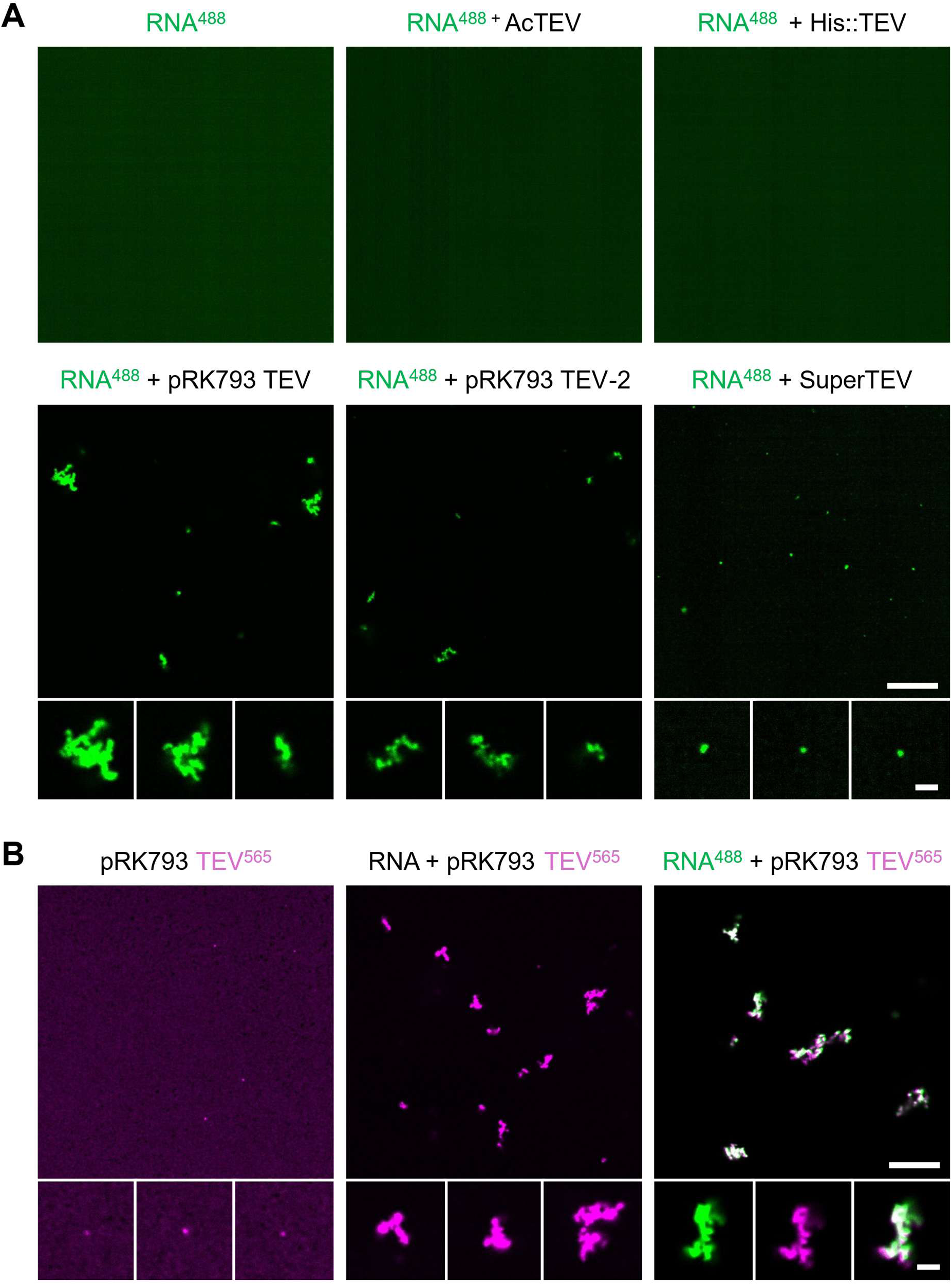
TEV proteases aggregate with RNA. A) Fluorescence images of 20 ng/µL fluorescently labeled *nos-2* RNA-Alexa-488 (green) when incubated with indicated TEV protease (0.4 U/µL AcTEV or 2.5 µM His::TEV, pRK793 TEV, pRK793 TEV-2, or SuperTEV). Scale bars: 20 µm, inset: 2 µm. Insets represent 3 distinct aggregates. B) Images of pRK793 TEV protease trace labeled with ATTO-565 (pink) incubated with and without labeled and unlabeled *nos-2* RNA as indicated (green). Scale bars: 20 µm, inset: 2 µm. Left and middle panels: insets represent 3 distinct aggregates. Right panel: insets represent individual channels of single aggregate.

We next asked whether pRK793 TEV protease itself aggregates under these conditions. Fluorescently trace-labeled pRK793 TEV protease was uniformly distributed in the absence of RNA, indicating that it does not form aggregates on its own. In contrast, addition of unlabeled RNA resulted in aggregation of fluorescently labeled pRK793 TEV protease (Fig. 1B). When fluorescently labeled RNA and fluorescently labeled pRK793 TEV protease were incubated together, both signals colocalized within the same aggregates (Fig. 1B), confirming that RNA and TEV protease form co-aggregates.

### RNA aggregation is not dependent on fluorescent labeling

To investigate whether fluorescent dyes influenced aggregation (Ng et al. 2021), we employed two alternative methods to detect aggregates, including those potentially unobservable by confocal microscopy: turbidity and mass photometry (MP). Turbidity assays identify the formation of large particles by measuring increased light scattering. As anticipated, neither RNA nor TEV protease alone increased turbidity over background (Fig. 2A). However, mixing pRK793 TEV protease with unlabeled RNA resulted in increased turbidity, indicating aggregate formation (Fig. 2A). This increase was not observed when AcTEV was mixed with RNA, confirming microscopy results (Fig. 2A).

**Figure 2:**
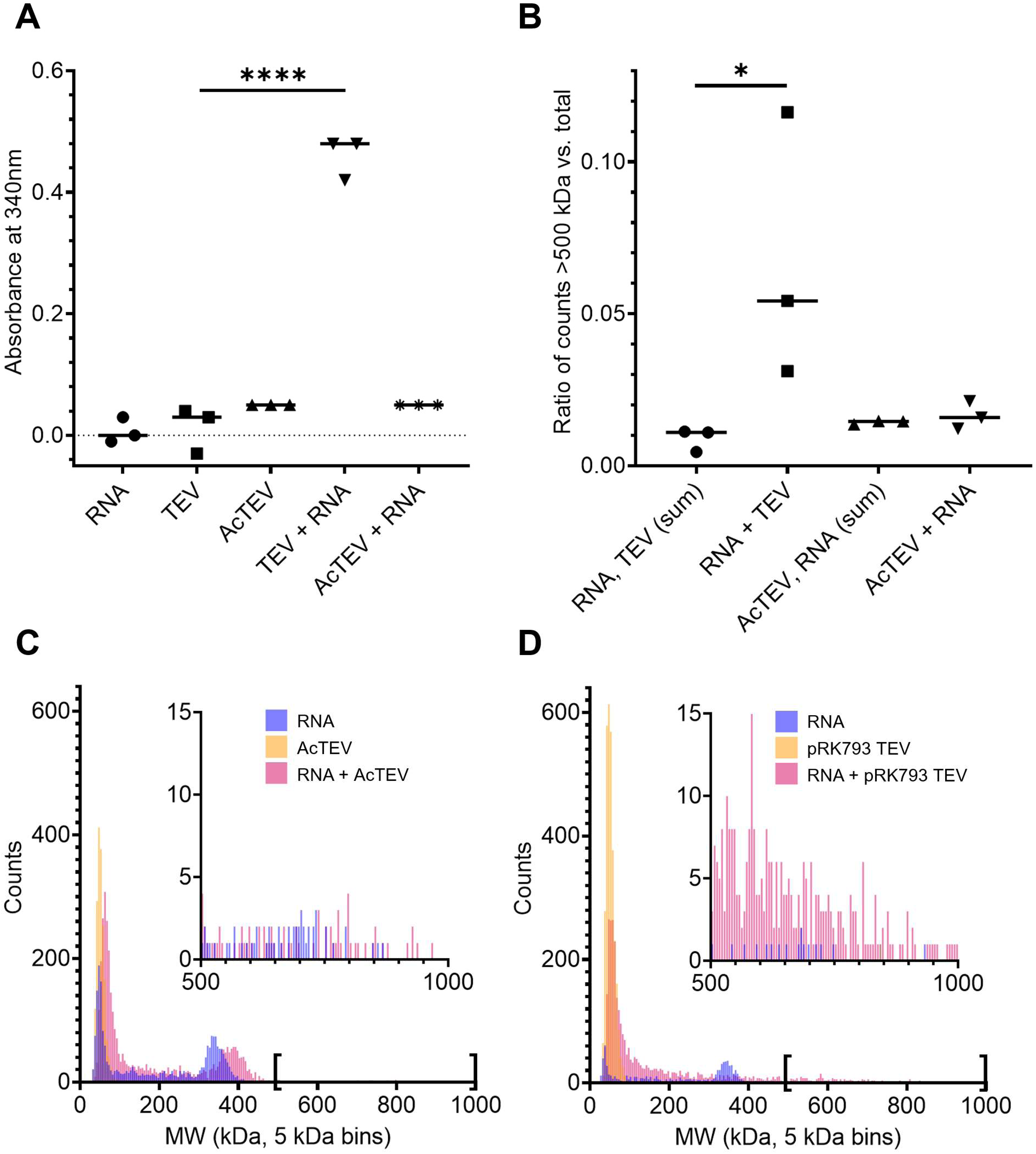
Detection of TEV protease–RNA complexes by mass photometry and turbidity. Representative mass photometry histograms obtained for 20 ng/µL *nos-2* RNA (blue), 2.5 µM pRK793 TEV (A) or 0.4 U/µL AcTEV protease (B) (orange), and TEV proteases incubated with *nos-2* RNA (pink) (5 kDA bin size). Insets show counts from 500-1000 kDa. (C) Ratio of particle counts larger than 500 kDa over total particle count for each condition indicated in A. Each point is an independent replicate. Lines represent the mean. (D) Turbidity was measured by absorbance at 340 nm of *nos-2* RNA with and without pRK793 TEV or AcTEV protease. Each point is an independent replicate. Lines represent the mean. One-way ANOVA test was used, *p<0.05, ****p<0.0001.

We next used MP to further characterize interactions between RNA and TEV protease (Fig. 2B-D). When analyzed alone, pRK793 TEV and AcTEV protease produced ratiometric contrast distributions corresponding to estimated molecular weights of approximately 53 kDa and 46 kDa, respectively (Fig. 2C and Fig. 2D). These values are consistent with the expected molecular weight of TEV protease (∼28 kDa), given that the practical lower detection limit of MP is approximately 30-50 kDa, which can lead to upwardly skewed apparent masses for species near this threshold (Kratochvíl et al. 2025 Oct 13). In MP measurements of RNA alone, a prominent peak was detected at approximately 344 kDa (Fig. 2C and Fig. 2D), consistent with the expected mass of the *nos-2* transcript. We also observed a recurrent peak near 42 kDa, which has been reported previously for long RNAs and is thought to arise from RNA molecules that do not adhere to the glass surface required for MP detection (Li et al. 2020; Schmudlach et al. 2025).

Upon mixing pRK793 TEV protease with RNA, we observed a loss of the free RNA peak at 344 kDa (Fig. 2D). Concomitantly, there was a significant increase in higher-molecular weight species exceeding 500 kDa, consistent with formation of large RNA-TEV assemblies (Fig. 2B and Fig. 2D). In contrast, incubation of AcTEV with RNA did not result in the appearance of these higher-molecular weight species (Fig. 2B and Fig. 2C), indicating that AcTEV does not undergo extensive aggregation with RNA under these conditions. Notably, however, we detected a reproducible shift of approximately 22 kDa in the apparent molecular weight of the RNA peak upon addition of AcTEV (Fig. 2C), indicating that AcTEV may associate with RNA without forming large aggregates.

### TEV protease-RNA aggregation salt dependence

We next investigated the effect of monovalent salt concentration on RNA-TEV aggregation, as both protein aggregation and protein-RNA interactions are sensitive to ionic strength. Fluorescently labeled RNA was incubated with either pRK793 TEV protease or AcTEV protease in buffers containing 50, 150, or 250 mM NaCl and imaged by confocal microscopy. AcTEV protease did not induce detectable RNA aggregation at 150 or 250 mM NaCl, but limited RNA aggregation was observed at 50 mM NaCl (Fig. 3A). For pRK793 TEV protease, RNA aggregates were most prevalent at 150 mM NaCl, reduced at 50 mM NaCl, and rare at 250 mM NaCl (Fig. 3A), though still more abundant than RNA alone under the same conditions. These results indicate that RNA-TEV aggregation is dependent on reaction conditions and that even TEV protease variants which are largely soluble under one set of conditions may exhibit aggregation at altered ionic strength.

**Figure 3:**
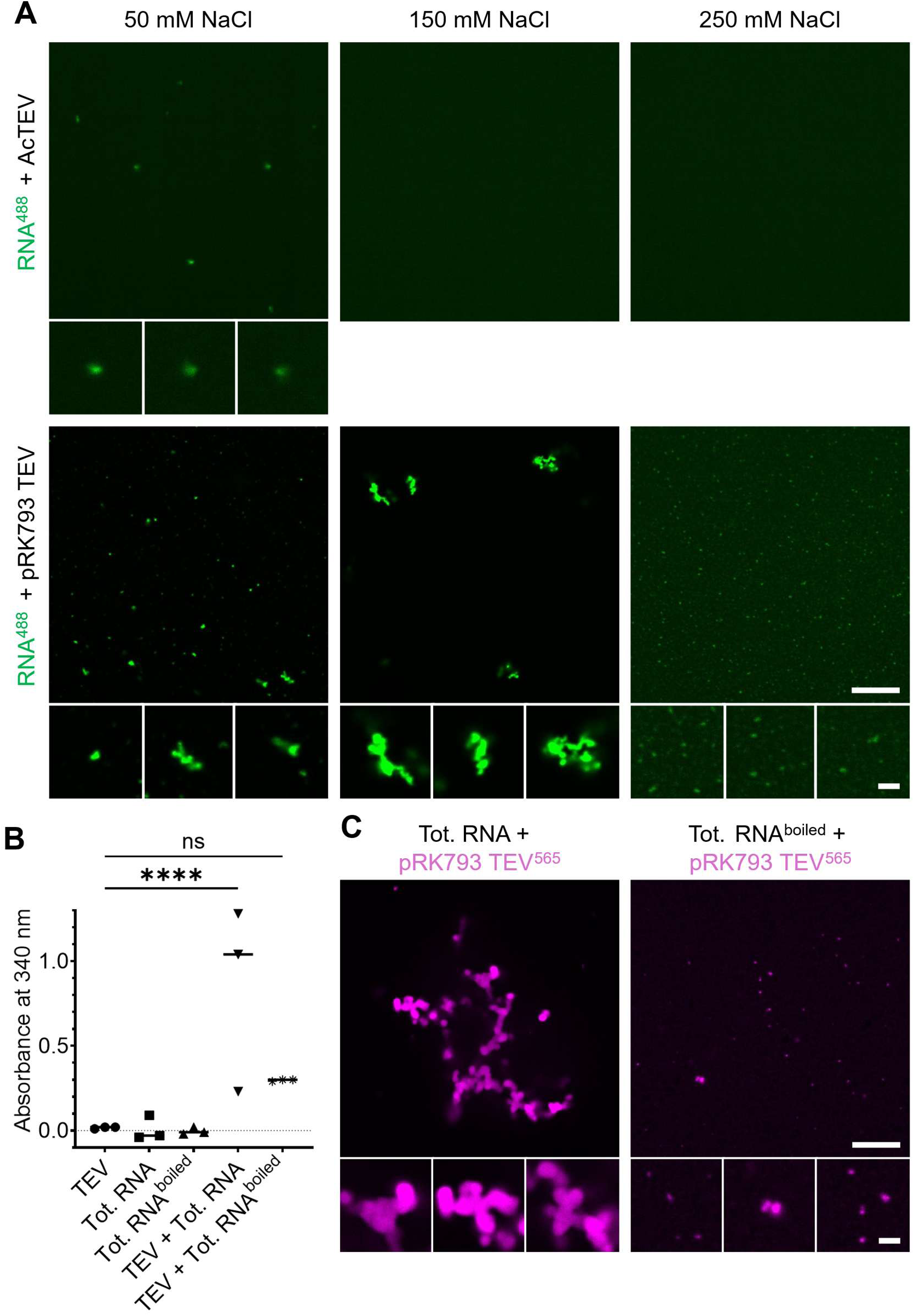
Salt concentration and RNA substrate influence TEV protease aggregation. (A) Images of 20 ng/µL *nos-2* RNA-Alexa-488 (green) with 2.5 µM pRK793 TEV or 0.4 U/µL AcTEV protease at 50, 150, and 250 mM NaCl. Scale bars: 20 µm, inset: 2 µm. (B) Turbidity was measured by absorbance at 340 nm of boiled and unboiled total RNA extracts from *C. elegans* with and without pRK793 TEV protease. (C) Images of fluorescently labeled pRK793 TEV protease-Alexa-488 (pink) with boiled and unboiled total RNA extracts from *C. elegans*. Scale bars: 20 µm, inset: 2 µm. One-way ANOVA test was used, ****p<0.0001, ns indicates no significance.

### TEV aggregation with other nucleic acids

To determine whether other nucleic acids induce similar aggregation with pRK793 TEV protease, we measured turbidity in reactions containing unlabeled pRK793 TEV protease. Incubation with total RNA isolated from *C. elegans* resulted in a marked increase in turbidity, consistent with aggregate formation (Fig. 3B). Boiling the total RNA prior to incubation reduced the turbidity signal, although levels remained above background (Fig. 3B), suggesting that RNA integrity or higher-order structure contributes to aggregation.

To confirm these observations, we visualized fluorescently labeled pRK793 TEV protease in the presence of the same nucleic acids by confocal microscopy. Incubation with total *C. elegans* RNA resulted in the formation of large pRK793 TEV protease aggregates (Fig. 3C). Aggregates were still observed when boiled total RNA was used, but they were fewer and smaller than those formed with untreated RNA (Fig. 3C), consistent with the turbidity measurements. We next tested whether other nucleic acids could induce aggregation detectable by confocal microscopy. No large aggregates of fluorescently labeled pRK793 TEV protease were observed in the presence of pUC19 plasmid DNA, 80 nt ssDNA, 21 nt ssRNA, or 21 nt ssDNA (Supplemental Table 2 and Supplemental Fig. 1), although small puncta were detectable at higher levels than in the absence of TEV protease. Together, these results suggest that pRK793 TEV protease aggregation is RNA-dependent and may be impacted by RNA length or higher-order RNA structure.

### Condensation competition results

We next investigated whether aggregation of TEV protease influences protein-RNA phase separation *in vitro*. We first examined PGL-3, an RNA-binding protein that undergoes salt-dependent LLPS and forms RNA-associated condensates that can be visualized by confocal microscopy (Saha et al. 2016). To test whether RNA-TEV aggregates alter PGL-3 condensate formation, we imaged fluorescently labeled PGL-3 and fluorescently labeled RNA in the presence or absence of pRK793 TEV protease or AcTEV protease and assessed changes in condensate morphology.

In the presence of AcTEV protease, PGL-3 and RNA formed condensates that were indistinguishable from those observed in the absence of protease (Fig. 4A). In contrast, addition of pRK793 TEV protease produced concentration-dependent effects on PGL-3-RNA assemblies. At low PGL-3 concentrations, we observed enhanced demixing of RNA and PGL-3, with morphologies resembling the RNA-TEV aggregates observedin the absence of PGL-3 (Fig. 1B, 4A). As the concentration of PGL-3 increased, PGL-3 and RNA formed more spherical structures that more closely resembled condensates formed without TEV protease. However, even under these conditions, PGL-3 condensates contained concentrated RNA puncta that were not detected in reactions lacking pRK793 TEV protease or containing AcTEV protease (Fig. 4A).

**Figure 4:**
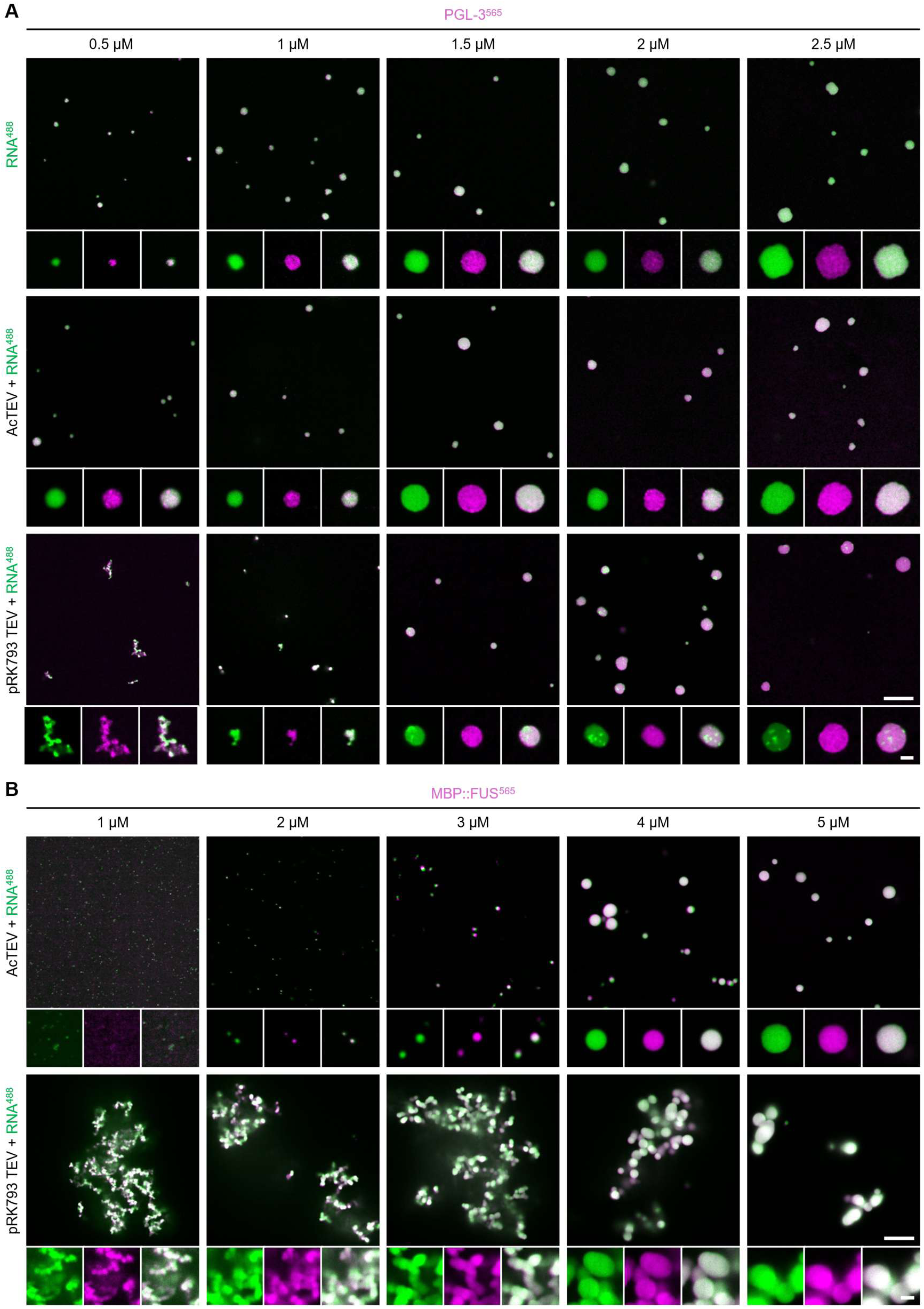
Aggregation of TEV protease and *nos-2* RNA influences condensate morphology in *in vitro* LLPS experiments. (A) Fluorescence images of a titration from 1-2.5 µM PGL-3 trace labeled with ATTO-565 (pink) incubated with fluorescently labeled 20 ng/µL *nos-2* RNA-Alexa-488 (green) in the presence or absence of 2.5 µM pRK793 TEV or 0.4 U/µL AcTEV protease. Scale bars: 20 µm, inset: 2 µm. (B) Images of a titration from 1-5 µM MBP::FUS trace labeled with ATTO-565 (pink) incubated with fluorescently labeled *nos-2* RNA-Alexa-488 (green) in the presence of pRK793 TEV or AcTEV protease. Scale bars: 20 µm, inset: 2 µm.

We next examined whether similar effects were observed with FUS, a well-characterized RNA-binding protein that undergoes RNA-dependent phase separation *in vitro* (Burke et al. 2015; Rhine et al. 2020). In this experimental system, TEV protease induces LLPS by cleaving an N-terminal maltose-binding protein (MBP) tag from FUS (Burke et al. 2015; Rhine et al. 2020). When 1 μM FUS was incubated with AcTEV protease, we observed small FUS-RNA assemblies (Fig. 4B). Increasing the concentration of FUS resulted in the formation of spherical FUS-RNA droplets, consistent with previous reports of liquid-like FUS condensates (Maharana et al. 2018; Rhine et al. 2020). In contrast, at 1 μM FUS incubated with pRK793 TEV protease, FUS and RNA colocalized within large, irregular aggregates (Fig. 4B). At higher FUS concentrations, FUS-RNA assemblies became more spherical but remained irregularly clustered, consistent with arrested coalescence relative to condensates formed in the presence of AcTEV protease.

Together, these results demonstrate that RNA-dependent aggregation of TEV protease can alter the morphology and organization of protein-RNA condensates and influence the apparent outcome of *in vitro* phase separation assays.

## Discussion

In this study, we demonstrate that TEV protease, a reagent used to induce LLPS *in vitro* through proteolytic removal of solubility tags, can undergo RNA-dependent aggregation and interact with RNA. Using confocal microscopy, turbidity measurements, and mass photometry, we show that the commonly used pRK793 TEV protease aggregates in the presence of long single-stranded RNA. Additionally, we find that distinct TEV protease variants exhibit distinct aggregation behaviors under identical conditions. Together, these results indicate that TEV protease is not universally inert in RNA-protein condensate assays and can alter condensate morphology, coalescence, and internal organization.

TEV protease is often assumed to be inert during condensate reconstitution, yet early biochemical work reported nonspecific TEV-RNA interactions and tag-dependent effects on solubility that have received limited attention in the context of biomolecular condensation (Daròs and Carrington 1997; Blommel and Fox 2007). Building on these observations, our microscopy and mass photometry data show that multiple TEV variants can associate with RNA, with the 5xArg tag markedly enhancing RNA binding and aggregation. The 5xArg extension was introduced to improve purification and solubility of the TEV catalytic core, but its arginine-rich composition likely promotes electrostatic interactions with RNA, highlighting a potential tradeoff between solubility and RNA association in RNA-rich LLPS assays. Consistent with this interpretation, addition of AcTEV protease resulted in a complete shift of the free *nos-2* RNA peak in mass photometry even in the absence of aggregation, indicative of RNA binding. While the molecular details of this interaction were not fully resolved here, these findings demonstrate that TEV protease variants can associate with RNA under commonly used condensate reconstitution conditions. These results will motivate future efforts to define the determinants of TEV-RNA interactions and to engineer variants with reduced RNA binding while retaining efficient proteolytic activity. Critically, we find that RNA-TEV aggregation can compete with or perturb normal droplet assembly in well-characterized protein-RNA phase separation systems.

In addition to direct RNA interactions, the concentration of TEV protease and the kinetics of tag cleavage represent an additional pathway through which protease-mediated condensation can influence phase behavior. Recent studies have shown that condensates formed through protease-mediated tag removal can exhibit significantly different phase behavior compared with those induced by rapid changes in pH, ionic strength, or dilution of low concentrations of denaturants such as urea (Van Lindt et al. 2021; Ge et al. 2025). To achieve rapid and complete cleavage and thereby trigger phase separation on short experimental timescales, TEV protease is used at relatively high concentrations. However, these conditions will also favor TEV self-association and RNA-dependent aggregation, particularly in RNA-rich environments, increasing the likelihood that the protease itself contributes to condensate formation. In contrast, slower cleavage regimes achieved through lower protease concentrations may reduce TEV aggregation but can introduce non-equilibrium effects by extending intermediate states in which partially cleaved or locally concentrated protein promotes nucleation-dependent assembly (Ge et al. 2025). Thus, both cleavage kinetics and protease aggregation can shape observed phase behavior through distinct but overlapping mechanisms.

These considerations highlight the value of alternative strategies for inducing phase separation. Approaches such as rapid pH jumps or the addition of low concentrations of denaturants that modulate protein solubility directly have been suggested to provide more reproducible control over phase behavior (Van Lindt et al. 2021; Ge et al. 2025). Such methods may be particularly useful when the goal is to compare phase behavior across conditions or in the investigation of condensate assembly dynamics and intermediates. Nevertheless, protease-mediated tag removal remains an important and often necessary strategy, especially for proteins that are difficult to purify in a soluble form without solubilizing tags.

When protease-mediated induction is required, careful experimental design is essential to distinguish intrinsic properties of the target protein from contributions arising from auxiliary reagents. Important considerations include explicitly reporting the identity of the TEV protease used, including all mutations and affinity or solubility tags, assessing the behavior of the protease alone under identical conditions, and comparing multiple protease variants with distinct engineering features. Finally, directly investigating recruitment of TEV protease to the condensate phase using microscopy or sedimentation-based assays may help determine whether the protease partitions into condensates. Alternative proteases may also be considered, although their own propensity to interact with RNA or undergo phase separation should be evaluated. Protease variants with positively charged tags such as polyarginine should be avoided in experiments containing RNA. Taken together, our results and other recent studies demonstrate that TEV protease can influence *in vitro* protein-RNA condensate assays in a context-dependent manner through a combination of RNA binding, aggregation, and cleavage-driven kinetic effects (Van Lindt et al. 2021; Ge et al. 2025). By accounting for these factors and carefully validating induction strategies, future studies can more accurately interpret condensate behavior and better isolate the biophysical principles governing phase separation.

## Methods

### pRK793 and SuperTEV purification

Plasmids containing the pRK793 TEV and SuperTEV protease were obtained from Addgene (#8827 and #193833, respectively). TEV proteases were expressed in Rosetta 2 (DE3) pLysS cells in 1 L terrific broth with ampicillin (100 µg mL^-1^) and chloramphenicol (100 µg mL^-1^) by growing at 37°C to an OD_600_ of approximately 1.0 and inducing overexpression with 1 mM IPTG at 16°C for 16 hours. Cells were resuspended in 50 mL Buffer A (20 mM HEPES pH 8.0, 500 mM NaCl, 25 mM imidazole, 10% (v/v) glycerol, 0.2 mM MgCl_2_, 2 mM DTT, 0.4 mM PMSF, 2 U/mL DNAse I (Sigma, cat. 4716728001), 10 µg/mL RNAse A (ThermoFisher, cat. EN0531), and protease inhibitors (ThermoFisher, cat. A32965)), lysed by microfluidization, and spun at 15,000g for 30 minutes at 4°C. TEV protease lysates were filtered and loaded onto a HisTrap HP 5 mL column (Cytiva, cat. 17524801) equilibrated with Buffer B (20 mM HEPES pH 8.0, 500 mM NaCl, 25 mM imidazole, 10% (v/v) glycerol), washed with Buffer B, and eluted with a gradient from Buffer B to Buffer B + 500 mM imidazole. SuperTEV protease received an additional step of purification by pooling peak fractions and loading onto a HiTrap Heparin HP 5 mL column (Cytiva, cat. 17040701) equilibrated with Buffer C (25 mM HEPES pH 7.5, 20% (v/v) glycerol, 2 mM DTT) + 100 mM NaCl while diluting 1/10-fold in Buffer C for an effective NaCl concentration of 50 mM. SuperTEV protease was washed and eluted with a gradient from Buffer C + 100 mM NaCl to Buffer C + 1 M NaCl. Peak fractions of pRK793 TEV or SuperTEV protease were pooled, concentrated to approximately 2 mL, and loaded onto a HiPrep 16/60 Sephacryl S-200 HR column (Cytiva, cat. 17116601) equilibrated with Buffer C + 500 mM NaCl and protein was eluted with Buffer C + 500 mM NaCl. Peak fractions were pooled, concentrated, aliquoted, flash frozen in liquid nitrogen, and stored at −80°C.

### pRK793 TEV-2 purification

pRK793 TEV-2 protease was expressed in Rosetta 2 (DE3) cells in 1 L LB broth with ampicillin (100 µg mL^-1^) by growing at 37°C to an OD_600_ of approximately 0.6 and inducing overexpression with 0.5 mM IPTG at 16°C for 16 hours. Cells were resuspended in 100 mL Buffer D (20 mM HEPES pH 7.5, 300 mM NaCl, 30 mM imidazole, 10% (v/v) glycerol, 2 mM DTT), lysed by microfluidization, and spun at 25,000g for 30 minutes at 4°C. 5% (v/v) PEI was added to a final concentration of 0.25% (v/v) over 25 minutes while stirring and allowed to incubate while stirring for an additional 15 minutes at 4°C. Lysate was clarified by centrifugation at 35,000g for 30 minutes at 4°C. Protein was precipitated by adding solid ammonium sulfate to a final concentration of 0.39 g/mL and allowed to incubate while stirring at 4°C for 16 hours. Precipitated protein was centrifuged at 25,000g for 30 minutes at 4°C, resuspended in 50 mL Buffer D, and centrifuged again at 25,000g for 30 minutes at 4°C. Supernatant was filtered and incubated with 2 mL Ni-NTA agarose beads (Qiagen, cat. 30210) equilibrated with Buffer D for 1 hour. Beads were loaded onto a gravity column, washed with Buffer D, and eluted with Buffer D + 300 mM imidazole. Eluent was concentrated to approximately 2 mL, loaded onto a Supradex 200 Increase 10/300 GL column (Cytiva, cat. 28990944) equilibrated with Buffer E (20 mM HEPES pH 7.5, 300 mM NaCl, 5% (v/v) glycerol, 2 mM DTT), and eluted with Buffer E. Peak fractions were pooled, concentrated, aliquoted, flash frozen in liquid nitrogen, and stored at −80°C. Purification protocol was adapted from a previous protocol.

### His::TEV purification

His::TEV protease was expressed in Rosetta 2 (DE3) pLysS cells in 2 L terrific broth with ampicillin (100 µg mL^-1^) and chloramphenicol (34 µg mL^-1^) by growing at 37°C to an OD_600_ of approximately 2.0, then at 16°C for 30 minutes, and inducing overexpression with 1 mM IPTG at 16°C for 16 hours. Cells were resuspended in 45 mL Buffer F (20 mM Tris pH 8.0, 500 mM NaCl, 25 mM imidazole, 10% glycerol, 1 mM TCEP, 10 µg/uL DNAse I (Sigma-Aldrich, cat. DN25), 0.2 mg/mL lysozyme, (MP Biomedicals, cat. 100831), 0.5x protease inhibitors (Calbiochem, cat. 539134)), lysed by sonication, and spun at 20,000g for 30 minutes at 4°C. Lysate was filtered and loaded flowed over 3 mL Ni-NTA agarose beads (Qiagen, cat. 30210) equilibrated with Buffer G (20 mM Tris pH 8.0, 500 mM NaCl, 25 mM imidazole, 10% glycerol, 1 mM TCEP), washed with Buffer G, and eluted with Buffer G + 500 mM imidazole. Peak fractions were pooled, concentrated, aliquoted, flash frozen in liquid nitrogen, and stored at −80°C.

### PGL-3 Purification

MBP::6xHis::TEV::PGL-3 was expressed in Rosetta 2 (DE3) pLysS cells in 1 L terrific broth with ampicillin (100 µg mL^-1^) and chloramphenicol (100 µg mL^-1^) by growing at 37°C to an OD_600_ of approximately 1.0 and inducing overexpression with 1 mM IPTG at 16°C for 16 hours. PGL-3 protein purification was adapted from a previous protocol (Putnam et al. 2019). Cells were resuspended 50 mL Buffer H (20 mM HEPES pH 7.5, 400 mM NaCl, 16% (v/v) glycerol, 0.4 M Arginine pH 8.0, 1% (v/v) IGEPAL, 0.2 mM MgCl_2_, 2 mM DTT, 0.4 mM PMSF, 2 U/mL DNase I, 10 µg/mL RNase A, and protease inhibitors), lysed by microfluidization, and spun at 15,000xg for 30 minutes at 4°C. Lysate was filtered and passed over 10 mL amylose resin (New England Biolabs, cat. E8021L) equilibrated with Buffer I (25 mM HEPES pH 7.5, 500 mM NaCl, 20% (v/v) glycerol, 2 mM DTT). Resin was washed with Buffer I, and protein was eluted with Buffer I + 20 mM maltose. Peak fractions were pooled and 1 mg of pRK793 TEV protease per 5 mg of protein was allowed to cleave for 16 hours at room temperature to remove the MBP tag. Solution was loaded onto a HisTrap HP 5 mL column equilibrated with Buffer B and washed with Buffer B. Flow-through was loaded onto a HiTrap Heparin HP 5 mL column equilibrated with Buffer C + 100 mM NaCl while diluting 10-fold in Buffer C for an effective NaCl concentration of 50 mM. Protein was washed and eluted with a gradient from Buffer C +100 mM NaCl to Buffer C + 1 M NaCl. Peak fractions were pooled, concentrated to approximately 2 mL, and loaded onto a HiPrep 16/60 Sephacryl S-200 HR column equilibrated with Buffer C + 500 mM NaCl, and protein was eluted with Buffer C + 500 mM NaCl. Peak fractions were pooled, concentrated, aliquoted, flash frozen in liquid nitrogen, and stored at −80°C.

### MBP::FUS Purification

6His::MBP::TEV::FUS was expressed in Rosetta 2 (DE3) pLysS cells in 1 L terrific broth with ampicillin (100 µg mL^-1^) and chloramphenicol (100 µg mL^-1^) by growing at 37°C to an OD_600_ of approximately 1.0 and inducing overexpression with 1 mM IPTG at 16°C for 16 hours. Cells were resuspended 50 mL Buffer H, lysed by microfluidization, and spun at 15,000xg for 30 minutes at 4°C. Lysate was filtered and passed over 10 mL amylose resin equilibrated with Buffer I. Resin was washed with Buffer I, and protein was eluted with Buffer I + 20 mM maltose. Peak fractions were pooled and loaded onto a HisTrap HP 5 mL column equilibrated with Buffer B, washed with Buffer B, and eluted with a gradient from Buffer B to Buffer B + 500 mM imidazole. Peak fractions were pooled and loaded onto a HiTrap Heparin HP 5 mL column equilibrated with Buffer C + 100 mM NaCl while diluting 10-fold in Buffer C for an effective NaCl concentration of 50 mM. Protein was washed and eluted with a gradient from Buffer C +100 mM NaCl to Buffer C + 1 M NaCl. Peak fractions were pooled, concentrated, aliquoted, flash frozen in liquid nitrogen, and stored at −80°C.

### Protein labeling

pRK793 TEV protease and PGL-3 were labeled with succinimidyl ester reactive fluorophores by mixing 70 µL protein at 25 µM in protein storage buffer with 5 µL ATTO-565 (ATTO-TEC, cat. AD 565) at 1 mg/mL in DMSO for 1 hour. Free fluorophore was removed by passage through three Zeba Spin desalting columns (7K MWCO, 0.5 mL) into protein storage buffer. The concentration of fluorophore-labeled protein and labeling efficiency were determined using a Nanodrop One spectrophotometer (ThermoFisher, cat. ND-ONE-W). Aliquots of labeled protein were flash frozen in liquid nitrogen and stored at −80°C.

### RNA transcription and labeling

*nos-2* RNA was transcribed using MEGAscript T7 Transcription Kit (ThermoFisher, cat. AM1333) and manufacturer’s recommended protocol. To generate fluorescently labeled *nos-2* RNA, 1 µL ChromaTide Alexa Fluor 488-5-UTP (ThermoFisher, cat. C11403) was added per 20 µL reaction resulting in a ratio of 1:150 labeled:unlabeled UTP. Template DNA for transcription reaction was obtained by PCR amplification from a plasmid containing the *nos-2* cDNA sequence (Lee et al. 2020). Free NTPs and protein were removed by LiCl_2_ precipitation per manufacturer’s protocol. RNAs were resuspended in water and stored at −80°C. The integrity of RNA products was verified by agarose gel electrophoresis (Supplemental Fig. 2A).

### TEV and oligonucleotide aggregation assays

TEV and oligonucleotide aggregation was induced by diluting TEV proteases from storage buffers to Buffer T (25 mM HEPES pH 7.5, 150 mM NaCl, 7% (v/v) glycerol, 0.6 mM DTT, 25 nM TCEP, 0.01% (v/v) IGEPAL, 0.004% (w/v) Triton X-100). 2.5 μM TEV protease was incubated with 20 ng/μL oligonucelotide (*nos-2* RNA*, nos-2* RNA^488^, 21 nt ssRNA (IDT), 21 nt ssDNA (IDT), 80 nt ssDNA (IDT), pUC19 (NEB, cat. N3041A), *C. elegans* total RNA, or *C. elegans* total RNA boiled at 95°C for 5 minutes prior to use) for 30 minutes at 20°C in Buffer T.

Total *C. elegans* RNA was prepared as described (He 2011). Fluorescently labeled pRK793 TEV protease was included as a trace label at a 1:100 labeled:unlabeled ratio as indicated. For salt titration assays, Buffer T contained 50, 150, or 250 mM NaCl as indicated. 5 µL of each solution were pipetted onto a coverslip, allowed to settle for 60 seconds, and imaged using a 40x-1.3NA objective with a 2.8x relay lens and illuminated with 488 and 565 nm lasers at an exposure time of 100ms. Experiments were performed in triplicate, and four images of each replicate were taken.

### Turbidity assays

Turbidity assays were performed by measuring absorbance at 340 nm on a Nanodrop One spectrophotometer. 2.5 μM unlabeled TEV or 0.4 U/µL AcTEV and 20 ng/μL unlabeled oligonucleotide (*nos-2* RNA, *C. elegans* total RNA, or *C. elegans* total RNA boiled at 95°C for 5 minutes prior to use) were incubated for 30 minutes in Buffer T at 20°C. 2 μL samples were measured in triplicate.

### Spinning disk confocal microscopy

Imaging of *in vitro* condensation assays were performed using a custom-built microscope (Intelligent Imaging Systems, 3i) inverted Zeiss Axio Observer with CSU-W1 SoRa spinning disk scan head (Yokogawa), 1X/2.8x/4x relay lens (Yokogawa), fast piezo z-drive (Applied Scientific Instrumentation), and an ORCA-Fusion BT Digital CMOS camera (Hamamatsu). Samples were illuminated with 488/561/637 nm solid-state laser (Coherent). Images were taken using Slidebook software using a 40x-1.3NA objective (Zeiss) and a 2.8x relay lens (Yokogawa) as indicated for each sample. All spinning disk confocal imaging was performed in a climate-controlled room at 20°C. See specific settings in sections below.

### Condensation assays

Protein condensation was induced by diluting proteins from storage buffers to final concentrations described in Buffer T. PGL-3 (0.5-2.5 μM as indicated) or MBP::FUS (1-5 μM as indicated) was incubated with 2.5 μM unlabeled TEV or 0.4 U/µL AcTEV and 20 ng/μL *nos-2* RNA^488^ for 30 or 120 minutes, respectively, at 20°C in Buffer T. Fluorescently labeled PGL-3 or MBP::FUS was included as a trace label at a 1:100 labeled:unlabeled ratio. 5 μL of each solution were pipetted onto a coverslip, allowed to settle for 60 seconds, and imaged using a 40x 1.3NA objective with a 2.8x relay lens and illuminated with 488 nm and 565 nm lasers at an exposure time of 100 ms. Experiments were performed in triplicate, and four images of each replicate were taken. Cleavage of MBP::FUS was verified via SDS-PAGE (Supplemental Fig. 2B).

### Mass photometry aggregation assays

Mass photometry (MP) assays were performed on a TwoMP mass photometer (Refeyn, cat. 02-0006). 2.5 μM unlabeled TEV or 0.4 U/µL AcTEV and 20 ng/μL unlabeled *nos-2* RNA were incubated for 30 minutes in Buffer T without 0.01% (v/v) IGEPAL at 20°C. Solutions were diluted 10-fold into Buffer T without 0.01% (v/v) IGEPAL, 5 μL of reaction was pipetted onto a glass coverslip coated with poly-L-lysine, and ratiometric contrast was measured for 60 seconds in triplicate. Apparent molecular weight (MW_app_) was determined using protein standards. The presence of 0.01% (v/v) IGEPAL in reactions generated high ratiometric noise and was omitted from MP assays.

## Acknowledgments

We would like to thank the Kirchdoerfer Lab at the University of Wisconsin-Madison for generously donating their preparation of pRK793 TEV-2 for this study. Research in the Putnam lab is supported by the National Institute of General Medical Sciences [R35 GM157010] and a David and Lucille Packard Fellowship for Science and Engineering. J.A.L. was supported in part by an NIH National Research Service Award [T32 GM158463].

## Author contributions

J.A.L. and A.A.P. conceived the overall project. The experimental plan and data collection were implemented by J.A.L and D.I.F., J.A.L. and A.A.P. prepared the figures and wrote the manuscript with suggestions from all authors. A.A.P. supervised all aspects of the work.

## References

1. Alberti S, Gladfelter A, Mittag T. 2019. Considerations and Challenges in Studying Liquid-Liquid Phase Separation and Biomolecular Condensates. Cell. 176(3):419–434. doi:10.1016/j.cell.2018.12.035.

2. Blommel PG, Fox BG. 2007. A combined approach to improving large-scale production of tobacco etch virus protease. Protein Expr Purif. 55(1):53–68. doi:10.1016/j.pep.2007.04.013.

3. Burke KA, Janke AM, Rhine CL, Fawzi NL. 2015. Residue-by-Residue View of In Vitro FUS Granules that Bind the C-Terminal Domain of RNA Polymerase II. Mol Cell. 60(2):231–241. doi:10.1016/j.molcel.2015.09.006.

4. Currie SL, Xing W, Muhlrad D, Decker CJ, Parker R, Rosen MK. 2023. Quantitative reconstitution of yeast RNA processing bodies. Proc Natl Acad Sci USA. 120(14):e2214064120. doi:10.1073/pnas.2214064120.

5. Daròs JA, Carrington JC. 1997. RNA binding activity of NIa proteinase of tobacco etch potyvirus. Virology. 237(2):327–336. doi:10.1006/viro.1997.8802.

6. Folkmann AW, Putnam A, Lee CF, Seydoux G. 2021. Regulation of biomolecular condensates by interfacial protein clusters. Science. 373(6560):1218–1224. doi:10.1126/science.abg7071.

7. Ge Y, Paul T, Gordiychuk M, Das N, Zhang Y, Myong S. 2025. FUS nanoclusters are a distinct state within the dilute phase. Nat Commun. 16(1):9956. doi:10.1038/s41467-025-64909-7.

8. He F. 2011. Total RNA Extraction from C. elegans. BIO-PROTOCOL. 1(6). doi:10.21769/BioProtoc.47. [accessed 2026 Feb 3]. https://bio-protocol.org/e47.

9. Kapust RB, Tözsér J, Fox JD, Anderson DE, Cherry S, Copeland TD, Waugh DS. 2001. Tobacco etch virus protease: mechanism of autolysis and rational design of stable mutants with wild-type catalytic proficiency. Protein Engineering, Design and Selection. 14(12):993–1000. doi:10.1093/protein/14.12.993.

10. Kratochvíl J, Van Wee R, Thiele JC, Loewenthal D, Bardzil J, Iqbal K, Benesch JLP, Thorpe S, Kukura P. 2025 Oct 13. Best practice mass photometry: a guide to optimal single-molecule mass measurement. Nat Protoc. doi:10.1038/s41596-025-01255-4. [accessed 2026 Feb 3]. https://www.nature.com/articles/s41596-025-01255-4.

11. Lau K, Bouchri B, Pojer F. 2023. Purification of 10xHis-SuperTEV v1. doi:10.17504/protocols.io.yxmvm2zxng3p/v1. [accessed 2026 Feb 3]. https://www.protocols.io/view/purification-of-10xhis-supertev-cn4uvgww.

12. Lee CYS, Putnam A, Lu T, He SX, Ouyang JPT, Seydoux G. 2020. Recruitment of mRNAs to P granules by condensation with intrinsically-disordered proteins. eLife. 9:e52896. doi:10.7554/eLife.52896.

13. Li Y, Struwe WB, Kukura P. 2020. Single molecule mass photometry of nucleic acids. Nucleic Acids Research. 48(17):e97–e97. doi:10.1093/nar/gkaa632.

14. Lin Y, Protter DSW, Rosen MK, Parker R. 2015. Formation and Maturation of Phase-Separated Liquid Droplets by RNA-Binding Proteins. Mol Cell. 60(2):208–219. doi:10.1016/j.molcel.2015.08.018.

15. Maharana S, Wang J, Papadopoulos DK, Richter D, Pozniakovsky A, Poser I, Bickle M, Rizk S, Guillén-Boixet J, Franzmann TM, et al. 2018. RNA buffers the phase separation behavior of prion-like RNA binding proteins. Science. 360(6391):918–921. doi:10.1126/science.aar7366.

16. Nautiyal K, Kuroda Y. 2018. A SEP tag enhances the expression, solubility and yield of recombinant TEV protease without altering its activity. New Biotechnology. 42:77–84. doi:10.1016/j.nbt.2018.02.006.

17. Ng SC, Güttler T, Görlich D. 2021. Recapitulation of selective nuclear import and export with a perfectly repeated 12mer GLFG peptide. Nat Commun. 12(1):4047. doi:10.1038/s41467-021-24292-5.

18. Parks TD, Leuther KK, Howard ED, Johnston SA, Dougherty WG. 1994. Release of Proteins and Peptides from Fusion Proteins Using a Recombinant Plant Virus Proteinase. Analytical Biochemistry. 216(2):413–417. doi:10.1006/abio.1994.1060.

19. Putnam A, Cassani M, Smith J, Seydoux G. 2019. A gel phase promotes condensation of liquid P granules in Caenorhabditis elegans embryos. Nature Structural & Molecular Biology. 26(3):220–226. doi:10.1038/s41594-019-0193-2.

20. Putnam A, Seydoux G. 2021. Cell-free reconstitution of multi-condensate assemblies. Methods Enzymol. 646:83–113. doi:10.1016/bs.mie.2020.07.004.

21. Rhine K, Makurath MA, Liu J, Skanchy S, Lopez C, Catalan KF, Ma Y, Fare CM, Shorter J, Ha T, et al. 2020. ALS/FTLD-Linked Mutations in FUS Glycine Residues Cause Accelerated Gelation and Reduced Interactions with Wild-Type FUS. Mol Cell. 80(4):666–681.e8. doi:10.1016/j.molcel.2020.10.014.

22. Saha S, Weber CA, Nousch M, Adame-Arana O, Hoege C, Hein MY, Osborne-Nishimura E, Mahamid J, Jahnel M, Jawerth L, et al. 2016. Polar Positioning of Phase-Separated Liquid Compartments in Cells Regulated by an mRNA Competition Mechanism. Cell. 166(6):1572–1584.e16. doi:10.1016/j.cell.2016.08.006.

23. Schmudlach A, Spear S, Hua Y, Fertier-Prizzon S, Kochling J. 2025. Mass photometry as a fast, facile characterization tool for direct measurement of mRNA length. Biol Methods Protoc. 10(1):bpaf021. doi:10.1093/biomethods/bpaf021.

24. Terpe K. 2003. Overview of tag protein fusions: from molecular and biochemical fundamentals to commercial systems. Appl Microbiol Biotechnol. 60(5):523–533. doi:10.1007/s00253-002-1158-6.

25. Tropea JE, Cherry S, Waugh DS. 2009. Expression and Purification of Soluble His6-Tagged TEV Protease. In: Doyle SA, editor. High Throughput Protein Expression and Purification. Vol. 498. Totowa, NJ: Humana Press. (Walker JM, editor. Methods in Molecular Biology). p. 297–307. [accessed 2026 Jan 28]. http://link.springer.com/10.1007/978-1-59745-196-3_19.

26. Van Lindt J, Bratek-Skicki A, Nguyen PN, Pakravan D, Durán-Armenta LF, Tantos A, Pancsa R, Van Den Bosch L, Maes D, Tompa P. 2021. A generic approach to study the kinetics of liquid–liquid phase separation under near-native conditions. Commun Biol. 4(1):77. doi:10.1038/s42003-020-01596-8.

27. Wadsworth GM, Srinivasan S, Lai LB, Datta M, Gopalan V, Banerjee PR. 2024. RNA-driven phase transitions in biomolecular condensates. Molecular Cell. 84(19):3692–3705. doi:10.1016/j.molcel.2024.09.005.

